# N-terminal 45 amino acids of DEF6 are necessary and sufficient to spontaneously colocalise with DCP1 in P-bodies

**DOI:** 10.1101/533109

**Authors:** Huaitao Cheng, Maha Alsayegh, Fred Sablitzky

## Abstract

DEF6 (Differentially Expressed in FDCP 6, also known as IBP and/or SLAT) is critical for the development of autoimmune disease and cancer. In T cells, DEF6 participates in TCR-mediated signalling determining T helper cell-mediated immune responses. In addition, DEF6 acts as a guanine nucleotide exchange factor (GEF) for Rho GTPases facilitating F-actin assembly and stabilisation of the immunological synapse (IS). However, DEF6 is also a component of mRNA processing bodies (P-bodies) linking it to mRNA metabolism. Including DEF6, more than 34 proteins have been shown to localise in P-bodies many of which contain a coiled coil domain, a super-secondary structure likely to facilitate interaction between these proteins. Accordingly, we suggested that the coiled coil domain in the C-terminal end of DEF6 was mediating P-body localisation of DEF6 under cellular stress conditions. However, a comprehensive analysis of wild type and mutant DEF6 proteins expressed in COS7 cells revealed that the coiled coil domain is dispensable for P-body colocalisation. Instead, we show here that the N-terminal 45 amino acids of DEF6 that contain one Ca2+-binding EF hand motif are sufficient to target DEF6 to P-bodies whereas the N-terminal 30 amino acids containing a disrupted EF hand motif are insufficient.

## INTRODUCTION

DEF6 (1), also known as SLAT (2) and IBP (3), was first isolated in a screen to identify differentially expressed genes during haematopoiesis (1). Subsequently, it was shown that DEF6 functions as a guanine nucleotide exchange factor (GEF) to activate Rho-GTPases (Rac1, Cdc42 and RhoA) (4, 5) coordinating actin cytoskeleton remodeling. DEF6 is highly expressed in T cells and upon T cell receptormediated activation, it is recruited to the junction between T cells and antigen presenting cells, termed immunological synapse (IS) (3, 6, 7). IS recruitment is dependent upon phosphorylation of DEF6 by LCK activating the GEF activity of DEF6 which in turn results in stabilisation of the IS (3, 6, 7). GEF function of DEF6 is also required for Ca2+-mediated NFAT signaling regulating Th1/Th2 differentiation and downstream inflammatory responses (6), In addition, DEF6 function seems critical in the development of autoimmune diseases by either promoting or inhibiting disease progress (7-13). We previously demonstrated DEF6 is also a substrate of the kinase ITK, phosphorylating Tyr210/222 of DEF6 in Jurkat T cells (14). DEF6 ITK phosphomimic mutant, Y210/222E, overexpressed in COS7 cells changed its cellular localisation forming DEF6 foci that overlapped with the P-body marker decapping enzyme 1 (DCP1). Endogenous DEF6 in Jurkat cells has also been observed to colocalise with P-bodies (14, 15). Moreover, arsenate or nocodazole treatment can induce WT DEF6 and several DEF6 mutants to colocalise with P-bodies in COS7 cells (14, 16). P-body is a major type of messenger ribonucleoproteins (mRNP) granule, which is non-membranous and dynamic cytoplasmic foci (17, 18). P-body is a supermolecular organelle that, including DEF6, contains more than 34 types of proteins (14, 19). Even though the components of P-bodies vary in different tissues, generally, the core of each P-body unit is mRNA decay machinery. The decay core can be simplified into two molecules, decapping enzyme 2 (DCP2) and XRN1. DCP2 responds to remove mRNA 5′-cap by catalysing the hydrolysis of the cap structure, and subsequently, XRN1, an exonuclease, degrades mRNA from 5′ to 3′. Even though DCP2 is capable to perform its function in vitro, its activity is stimulated and regulated by several other proteins termed decapping coactivators, which include DCP1, EDC3, EDC4, LSm1-7 complex, LSm14A, Dhh1 and Pat1. These proteins are highly conserved in eukaryotes, and all of which have been observed localised in P-bodies and function in 5′ to 3′ mRNA decay (20, for review see 21). In addition, small interfering RNAs (siRNA) induced RNA interference (RNAi) is also carried out in P-bodies by Argonaute-2 (Ago2), GW182 and/or the mRNA decay core (22). Moreover, mRNAs returning from P-bodies to polysome has also been observed in Yeast (23), which indicates P-bodies could also function as mRNA storage. DEF6 function in P-bodies is unkown and it remains to be seen which role DEF6 plays in this organelle. How P-bodies assemble is still not fully understood. Many factors are involved in this process, such as Ago, GW182 and mRNAs. This implies P-bodies formation is a complicated and multilevel process. Among them, EDC4 is a scaffold for the decay machinery (21). In P-bodies, EDC4 provides binding sites for DCP1, DCP2 and XRN1, where three DCP1 molecules gather as a trimer to activate DCP2 and subsequently performing the decapping. Without EDC4, DCP1 and DCP2 can still interact, however, the binding force between them is too weak to maintain their function. Moreover, the binding of DCP2 and XRN1 to the same EDC4 molecule ensures that the decapped mRNAs are transferred to XRN1 for further decay. Each of the decay machinery could be considered as a working unit. Assembly of these units to so-called microscopically visible P-body is also little understood. Decker and Parker (2012) summarised a possible model of P-body assembly in yeast, which depends on EDC3 self-interaction domain (Yjef-N) and Lsm4 Q/N rich domain (24). Although in *Drosophila* S2 cells, P-body assembly is not blocked by EDC3 depletion, depletion some P-body Q/N rich proteins, such as GW182, did decrease P-body formation in *Drosophila* and human cells. It is possible therefore that assembly through interaction of Q/N rich domains/proteins is the first step in P-body formation. Hence, a simple model of P-body formation can be summarised as following: in the first step, DCP1, DCP2, XRN1 rely on EDC4 as a scaffold to assemble as P-body units. Then these units gather to be microscopically visible P-bodies through the interactions of Q/N rich components. Q/N rich proteins tend to form coiled coil structure (25). DEF6 has a Q-rich coiled coil domain in its C terminal end. Thus, we proposed that phosphorylation of DEF6 liberates the coiled coil domain resulting in P-body colocalisation (14). Here, we report that the coiled coil domain of DEF6 is dispensable for DEF6 granule formation and colocalisation with DCP1 and that the N-terminal 45 amino acids of DEF6 are sufficient and necessary for spontaneous P-body localisation. The N-terminal 45 amino acids of DEF6 contain one Ca2+-binding EF hand motif (26) and disruption of which abolishes P-body localisation implying a potential relation between calcium signalling and mRNA metabolism.

## RESULTS

### The coiled coil domain of DEF6 is dispensable for granule formation and colocalisation with DCP1 in P-bodies

Hey et al. (2012) described that phosphorylation of DEF6 by ITK caused a conformational change resulting in DEF6 granule formation, colocalising with the P-body marker DCP1 (14). Given that P-body components often are proteins with coiled coil domains such as EDC4 and PAN3 (21, 17, 28), it was suggested that the conformational change released the coiled coil domain of DEF6 facilitating colocalisation and perhaps interaction with P-body components. The coiled coil motif is characterized by a heptad (seven) repeat denoted as (a-b-c-d-e-f-g)n, where the “a” and “d” positions are occupied by nonpolar amino acids, which provides the most significant force by mediating hydrophobic interactions with other “a” and “d” amino acids to stabilize the coiled coil structure (29, 30). To test the model proposed by Hey et al. (14), amino acids in ten “a” and “d” positions within the coiled coil domain of DEF6 were exchanged with proline residues. It had been shown that introduction of prolines disrupts the formation of coiled coil domains (31). Amino acid sequences alignments of DEF6 and its only related protein SWAP70 indicated that amino acids in the “a” and “d” positions are highly conserved across species from human to trichoplax (Suppl Fig. 1; 32). Based on these analyses, 10 a/d positions as indicated in Fig. 1A were selected and the amino acids exchanged with prolines using site-directed mutagenesis. Initially, 10 mutant DEF6 proteins with a single set of a/d mutations were established (Q371P-A374P, R407P-M410P, I428P-L431P, L442P-E445P, L463P-E466P, L470P-L473P, K491P-L494P, L505P-A508P, L512P-V515P and L533P-A536P), and subsequently, multiple sets of a/d mutations were combined with one DEF6 protein containing all 10 sets of a/d mutations (Q371P-A374P-L505P-A508P, R407P-M410P-L505P-A508P, I428P-L431P-L505P-A508P and All-10 mutant) (Fig. 1B). Sequence analysis of all mutants confirmed exchange of codons at the a/d positions chosen with no other apparent changes of the DEF6 coding region (Suppl. Fig. 4 and 5). COIL2 analysis shown in Fig. 1C suggested that introduction of proline residues indeed disrupts the coiled coil domain of DEF6. GFP-tagged DEF6 wild type and proline mutants were transfected into COS7 cells and the cellular localisation of the proteins was determined after 24h by fluorescent and/or confocal microscopy. COS7 cells do not express LCK nor ITK, two kinases known to phosphorylate DEF6 in T cells (14, 33) but it cannot be ruled out that other unknown modifications of DEF6 might occur. Nevertheless, changes in cellular localisation in non-treated COS7 cells are likely a consequence of conformational change due to the introduction of the proline residues rather than post-translational modifications. Compared to wild type DEF6 and GFP alone, that are diffuse in cytoplasm and in the case of GFP also in the nucleus, proline mutants exhibited three types of distinct phenotypes. R407P-M401P, I442P-E445P and K491P-L494P exhibited a diffuse localisation in the cytoplasm similar to wild type DEF6 (Fig. 2). Apart from Q371P-A374P, all other single set of a/d mutants as well as combined sets including the ALL-10 mutant seemed to mainly localise in filopodia, lamellipodia and podosomes suggesting colocalisation with F-actin. Q371P-A374P was mainly diffuse in the cytoplasm and somewhat localised with F-actin on lamellipodia but it also occasionally formed granules in the cytoplasm (Fig. 2). To test whether Q371P-A374P granules colocalise with P-bodies, a cotransfection with a P-bodies maker, DCP1 tagged with mCherry was carried out. Confocal analysis with Z-stack images shown in both vertical and side versions (X and Y coordinates) to provide a 3D view, revealed clearly that Q371-A374P granules did not colocalise with DCP1 (Suppl. Fig. 3) Arsenate treatment of cells results in an oxidative condition that leads to arrest of mRNA translation and induction of stress granules and P-bodies (34). It was previously shown that wild type DEF6 relocates and colocalises with DCP1 in P-bodies, after arsenate treatment (14). To test whether disruption of the coiled coil domain by the introduction of prolines would interfere with granule formation, COS7 cells were treated with 1mM arsenate for 40 mines 24 hrs after transfection. As shown in Fig. 3, wild type DEF6 formed granules in cytoplasm as expected. Apart from K491P-L494P mutant that exhibited granules in a subset of cells (17%), all other single set as well as combined sets including the All-10 mutant did not form granules. The data presented so far would suggest that the coiled coil domain is essential for granule formation of DEF6 in cells treated with arsenate. This is in line with data presented by Hey et al. (2012) who showed that DEF6 is phosphorylated by ITK at amino acid positions of Tyr 210 and Tyr 222 and that phosphomimc mutants spontaneously relocated colocalising with P-bodies. Therefore, the Tyr (Y) residues 210 and 222 were changed to Glu (E) in selected proline mutant proteins to test whether the disrupted coiled coil domain prevents spontaneous granule formation of the phosphomimic Y210-222E mutant. Mutant Y210-222E-Q371P-A374P, Y210-222E-I428P-L431P, Y210-222E-L505P-A508P, Y210-222E-Q371P-A374P-L505P-A508P, Y210-222E-I428P-L431P-L505P-A508P and Y210-222E-All-10 were established and transfected into COS7 cells. Surprisingly, all these combined mutants spontaneously formed granules in COS7 cells. Furthermore, cotransfections revealed that granules formed by the combined mutants colocalised with P-body marker DCP1 (Fig. 4). Confocal z-stack images also confirmed colocalisation of Y210-222E-L505P-A508P and Y210-22E-All-10 with DCP1 (Fig. 5). These results demonstrate that granule formation and P-body colocalisation of the phosphomimic Y210-222E mutant is independent of the coiled coil domain and suggests that instead another region of DEF6 is responsible for both.

**Figure 1.**
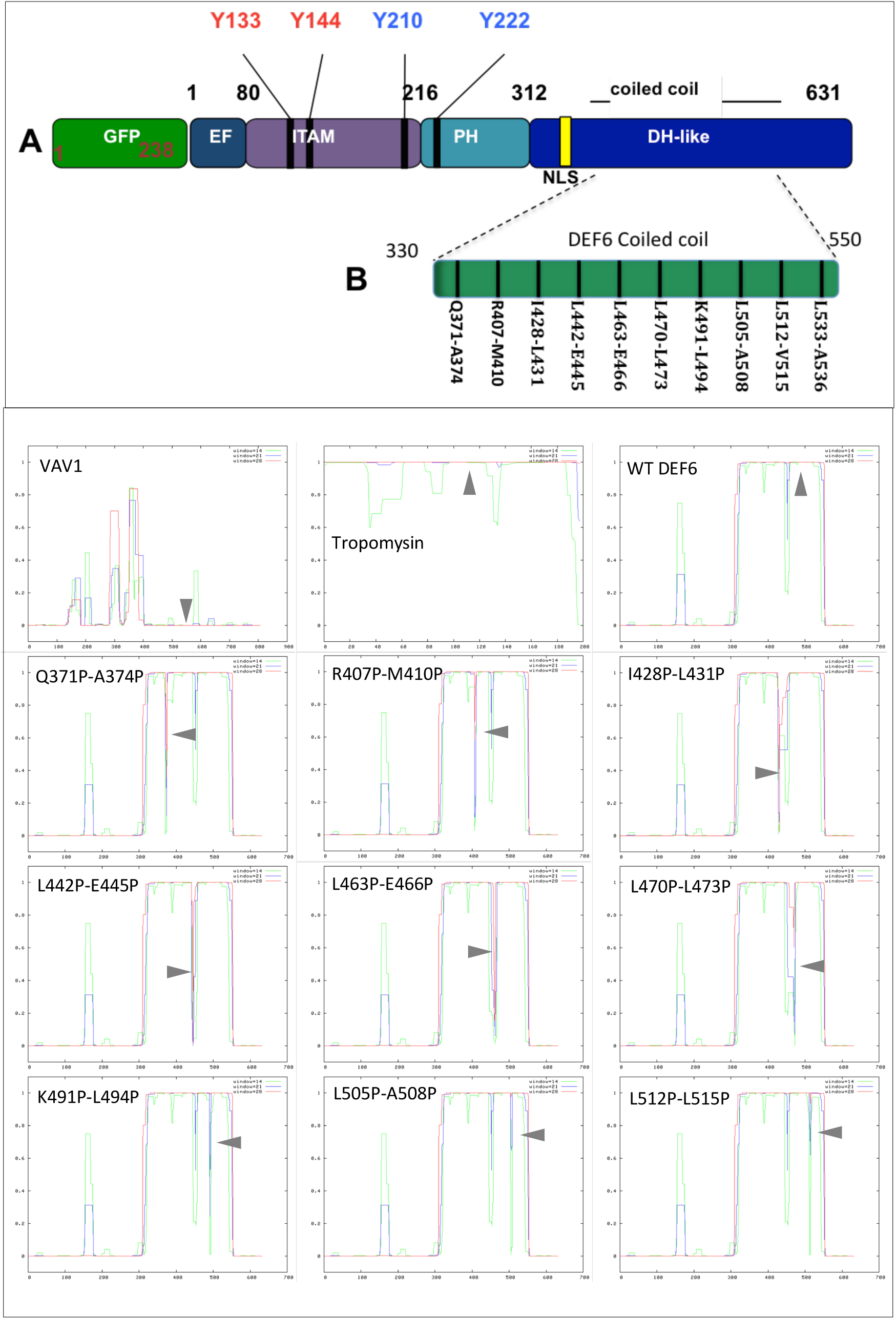
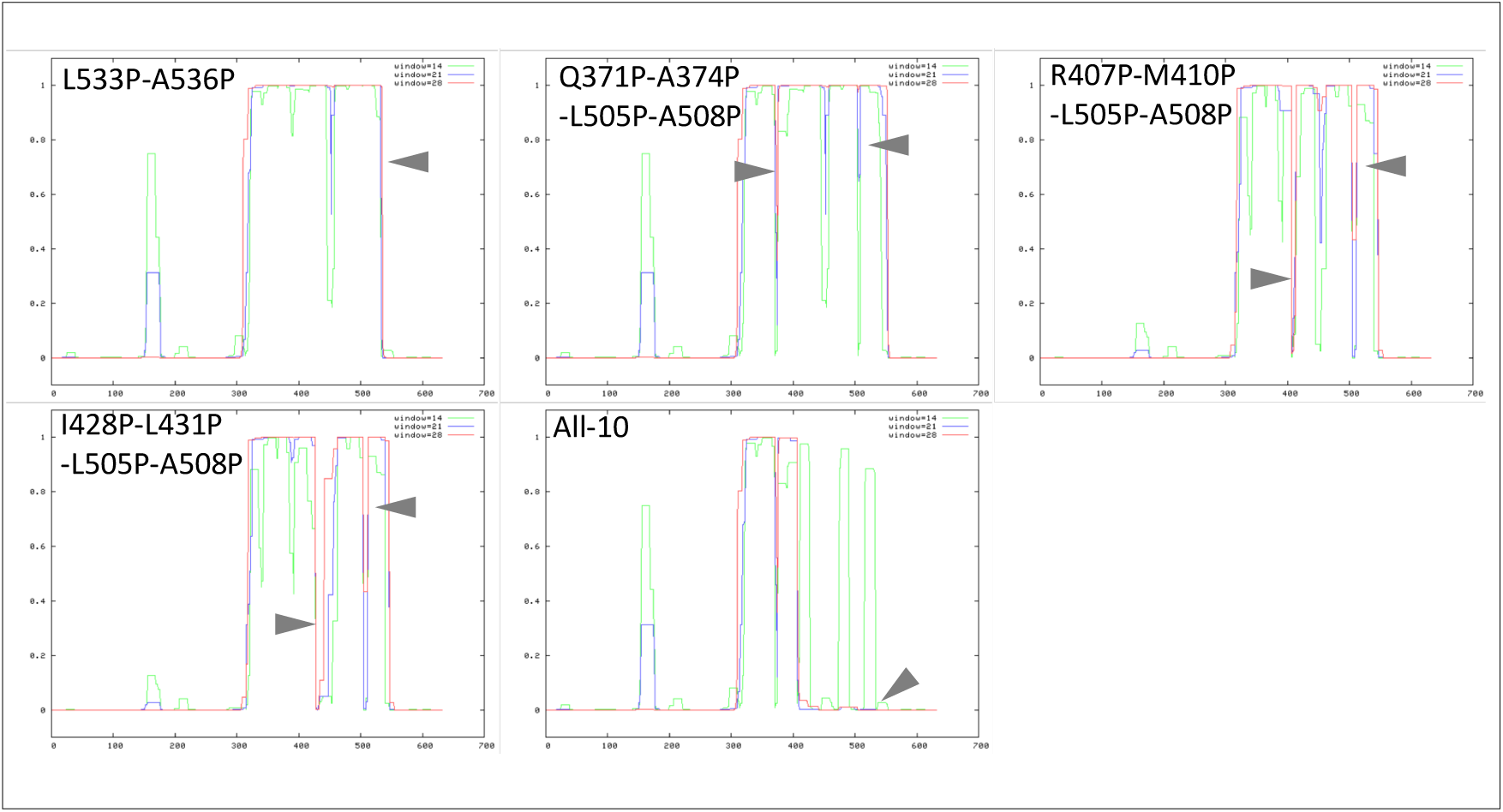
Insertion of proline residues at highly conserved a/d positions disrupts the coiled coil domain of DEF6 (A) Schematic representation of the domain structure of DEF6 fused to GFP. EF: Ca2+-binding EF-hand; ITAM: immunoreceptor tyrosine activation motifs within the DEF6-SWAP70 homology domain; PH: pleckstrin homology domain; DHL: Dbl homology like domain (see Figure 1.3.0). Within the ITAM and PH domains, Try-133/144 are phosphorylated by LCK and Try-210/222 by ITK. DHL domain contains a nuclear localisation sequence (NLS) and a coiled coil structure. (B) 10 highly conserved a/d positions within coiled coil sequence of DEF6 were selected and amino acids replaced by prolines (P) as indicated. (C) Coil2 analysis predicted disruption of the coiled coil domain in proline mutants compared to wild type DEF6. a/d double P mutants and ALL-10 predictions are shown as indicated. Vav1, lacking a coiled coil domain, served as a negative control and tropomysin that consists entirely out of a coiled coil structure as a positive control.

**Figure 2.**
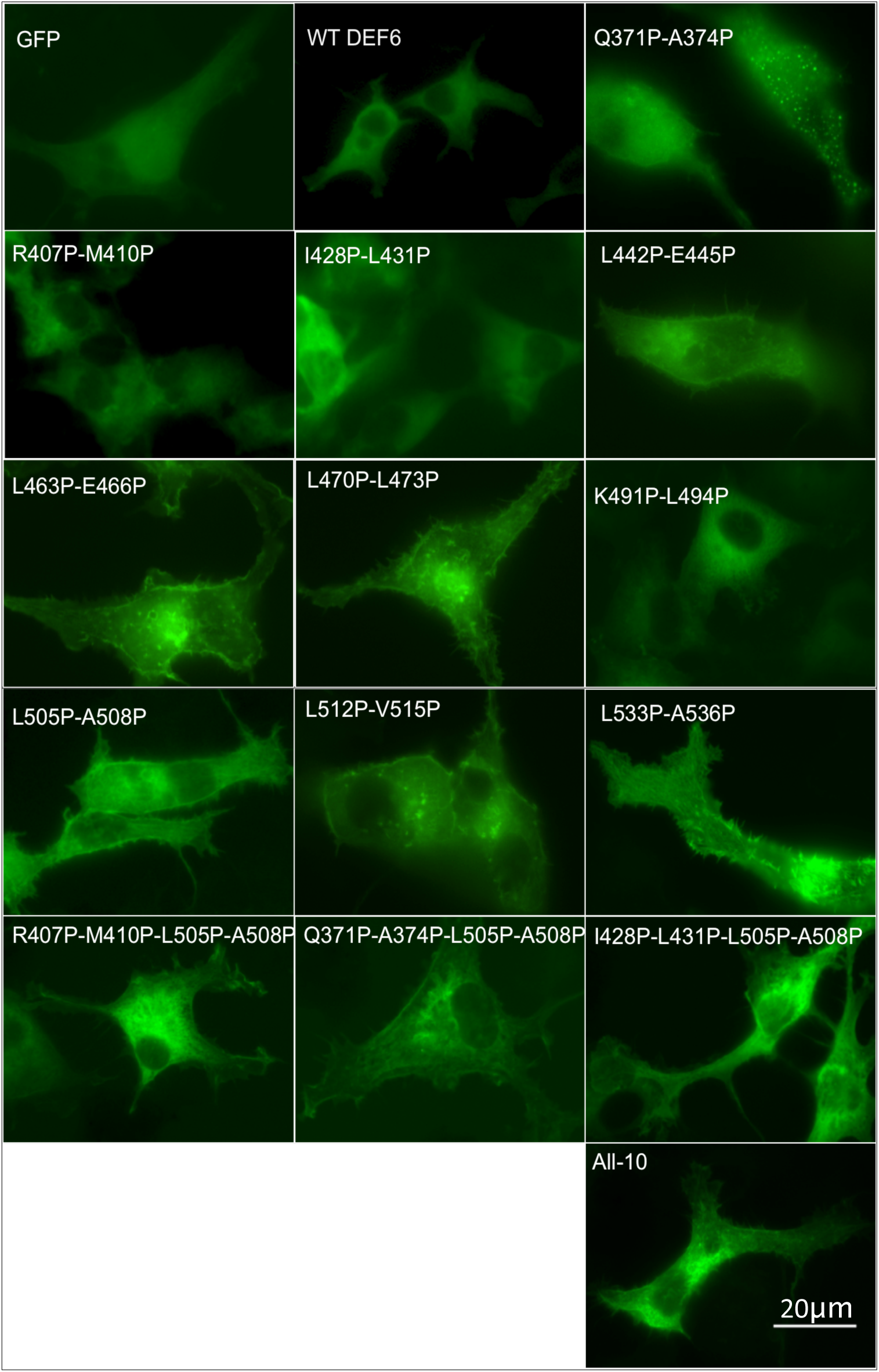
GFP-tagged DEF6 proline mutants exhibited three types of cellular localisation in COS7 cells. Type 1: Q371P-A374P was mainly diffuse in the cytoplasm but occasionally exhibited granules. Type 2: R407P-M410P, I428P-L431P and K491P-L494P were localised diffuse in the cytoplasm similar to WT DEF6. Type 3: All other mutant proteins as indicated localised in lamellipodia, filopodia and podosomes. GFP alone localised diffuse in the cytoplasm and nucleus.

**Figure 3.**
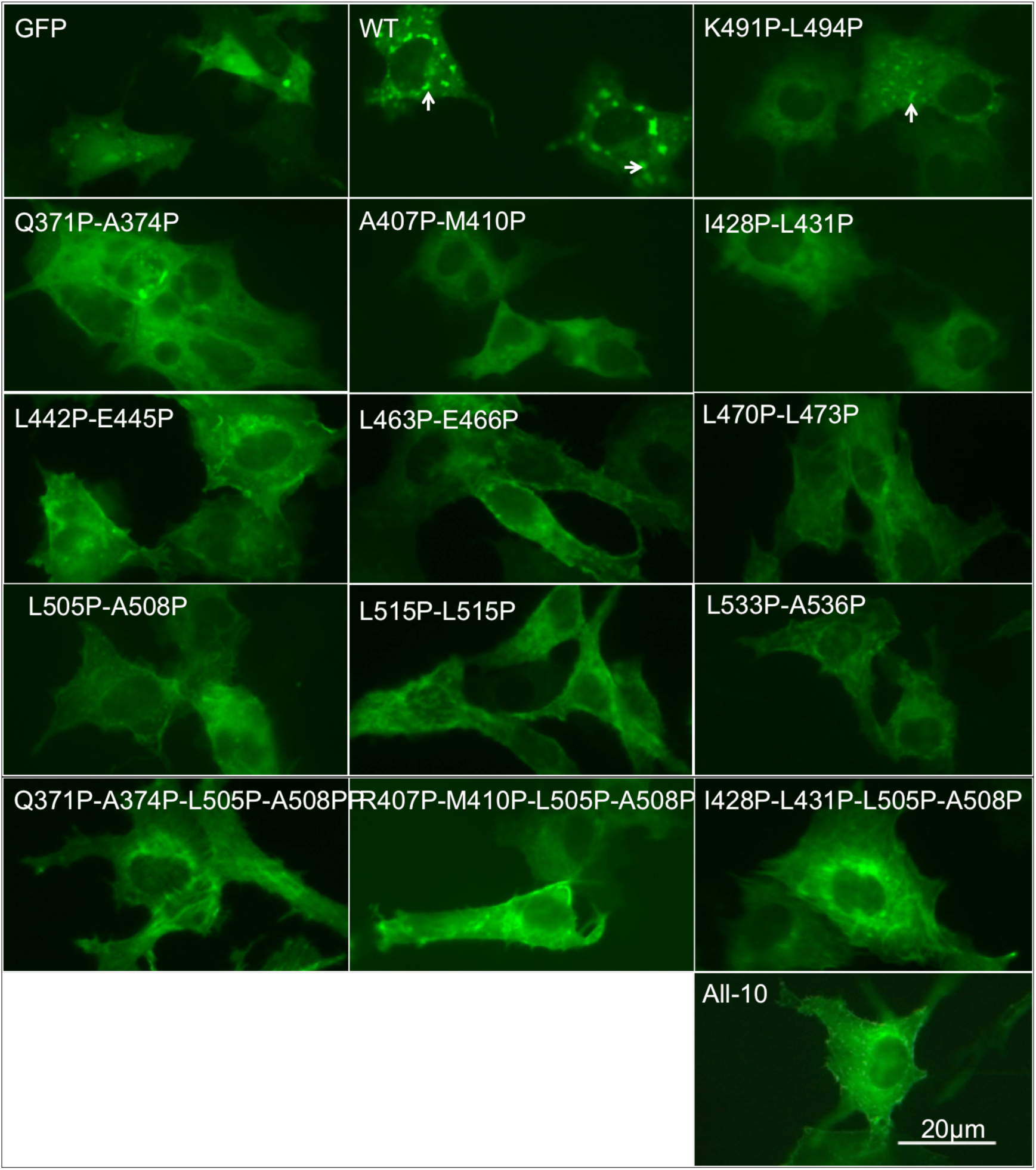
Disruption of the coiled coil domain prevented arsenate induced granule formation. Analysis of proline mutants as indicated 24 h after transfection of COS7 cells treated with arsenate (1mM, 40mins). As previously shown, WT DEF6 formed granules whereas the GFP control did not. Arsenate treatment had no effect on the proline mutants that exhibited similar cellular localisation as in untreated cells.

**Figure 4.**
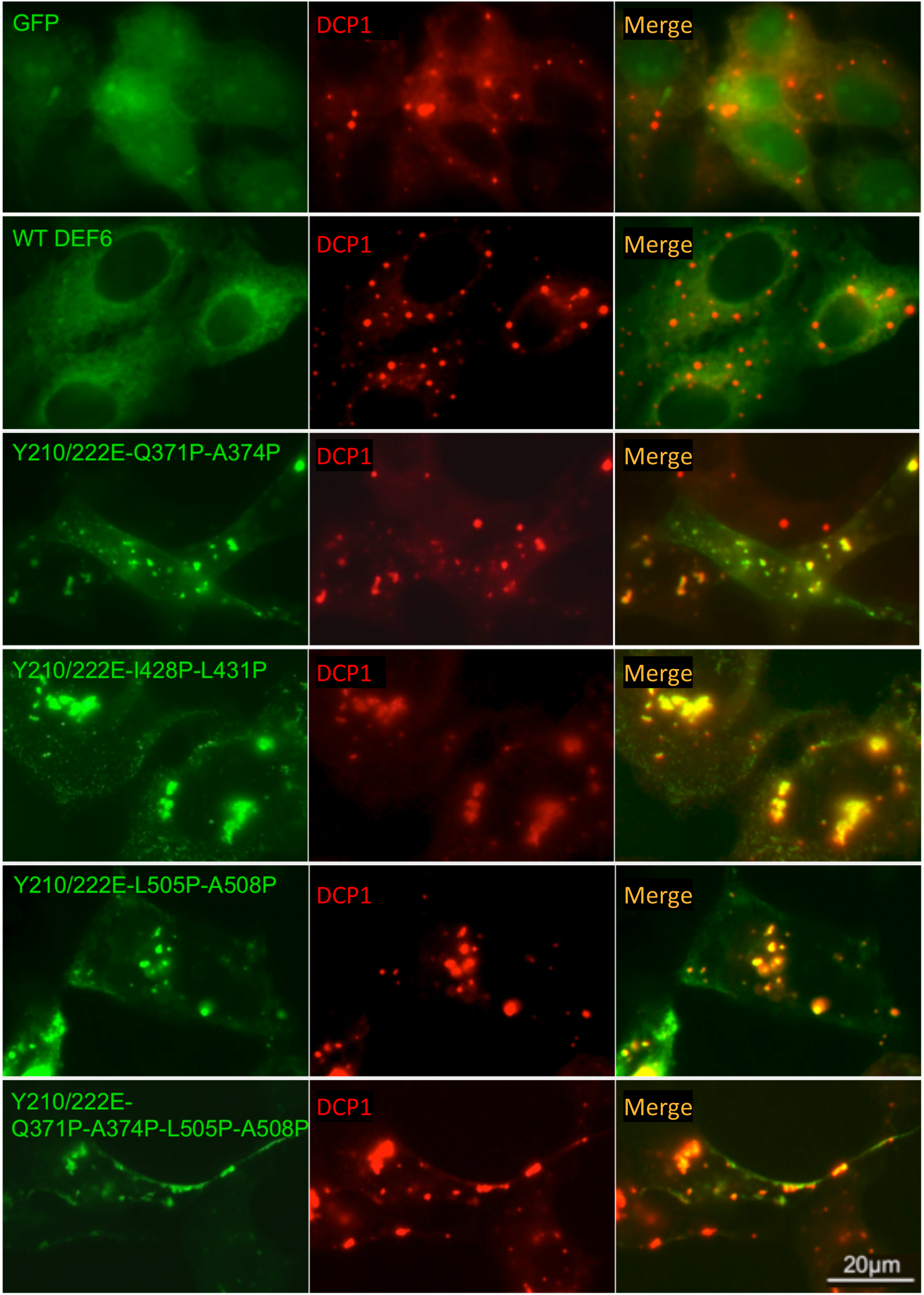
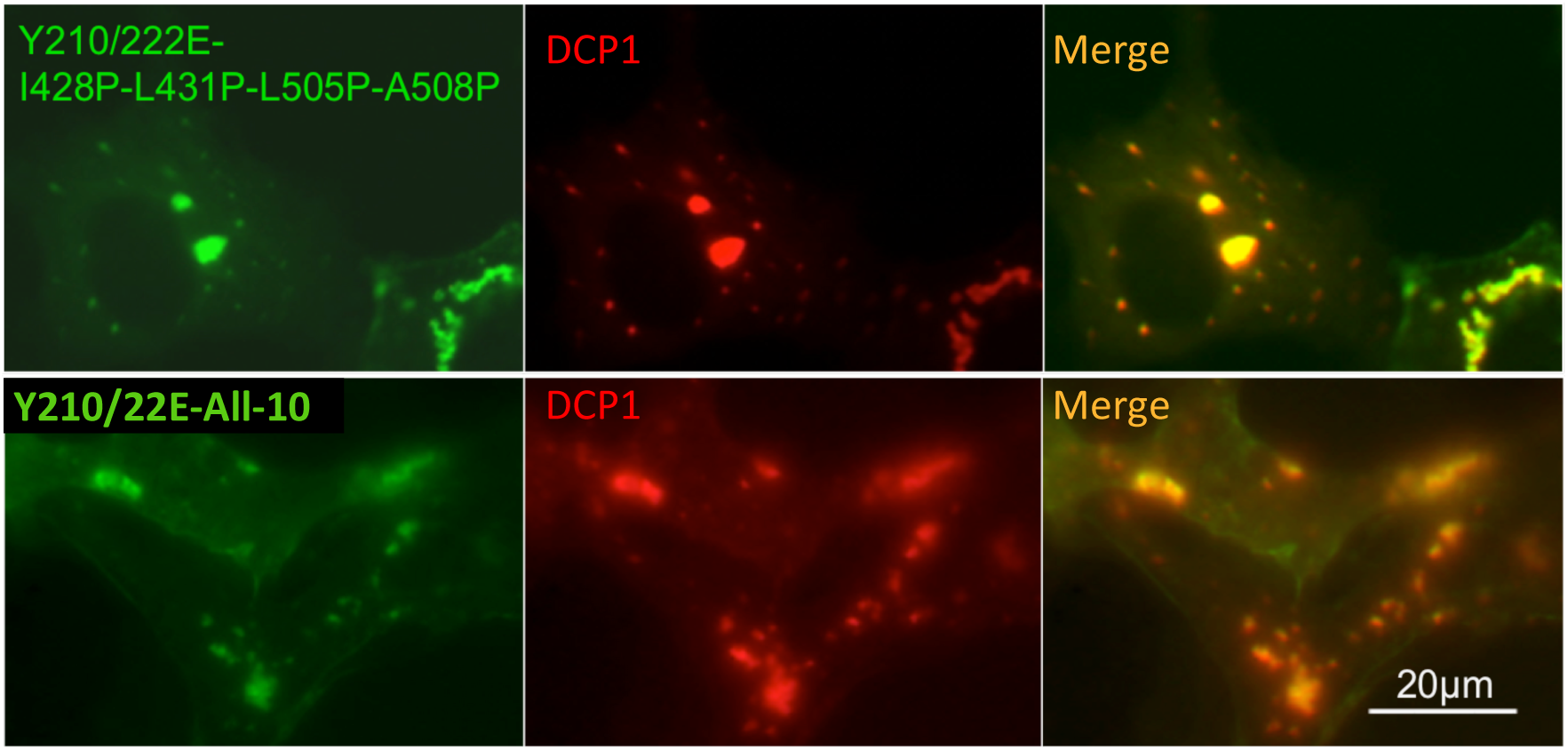
Introduction of the ITK phosphomimic mutations Y210/222E results in colocalisation of proline mutants with P-bodies. Y210/222E mutations were introduced into selected proline mutants as indicated and these combined GFP-tagged mutants (green) were cotransfected into COS7 with mCherry-tagged DCP1 (red). Images taken 24h after transfection. Merged images from the left and middle columns shown on the right indicate that the combined mutants formed granules that colocalised with DCP1.

**Figure 5.**
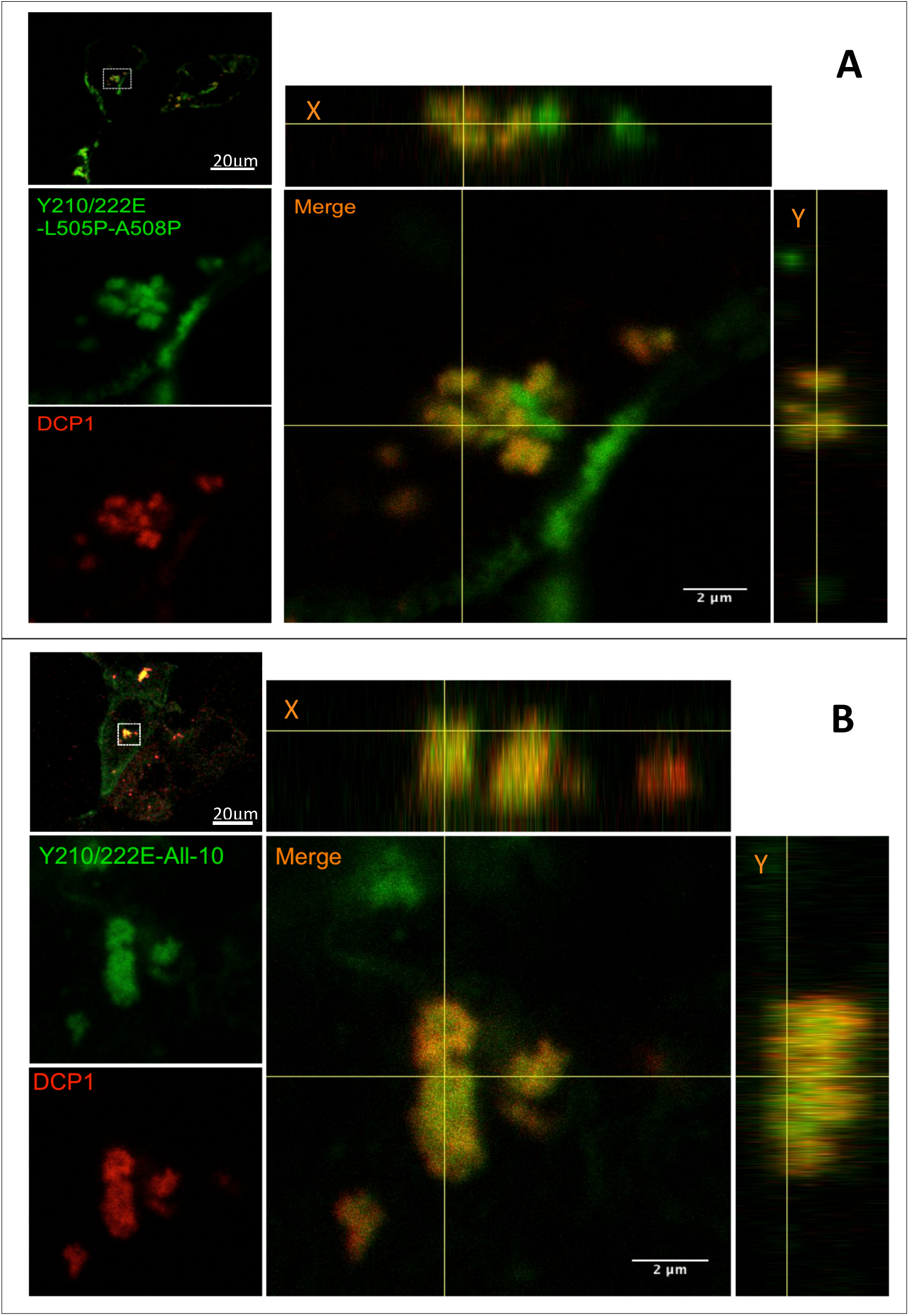
Y210/222E-L505P-A508P and Y210/222E-All-10 form granules that colocalise with DCP1. Confocal Z-stack analysis of GFP-tagged Y210/222E-L505P-A508P (A) and Y210/222E-All-10 (B) (green) cotransfected with mCherry-tagged DCP1 (red) 24 h after transfection of COS7 cells. Boxed area in the images on top left are enlarged on the left middle and bottom and further enlarged in the merged images on the right. Vertical and side views (X and Y coordinates) are also shown indicating colocalisation.

### N-terminal 45 amino acids of DEF6 are necessary and sufficient to spontaneously colo-calise with P-bodies

Having established that the coiled coil domain of DEF6 is dispensable for granule formation and colocalisation with P-bodies, C-terminal truncation mutants N-30, N-45, N-79, N-108, N-216 and N-312, N-590 were made as depicted in Fig. 6 and cellular localisation of these mutants tested in COS7 cells. All C-terminal truncation mutants but N-30 formed spontaneously granules (Fig. 6) and confocal Z-stack analysis indicated that N-45 granules completely overlapped with DCP1 (Fig. 7). Given that the EF hand motif in N-30 is disrupted (Suppl. Fig. 2), granule formation and colocalisation with DCP1 in P-bodies seems to depend on a functional Ca2+-binding EF hand motif.

**Figure 6.**
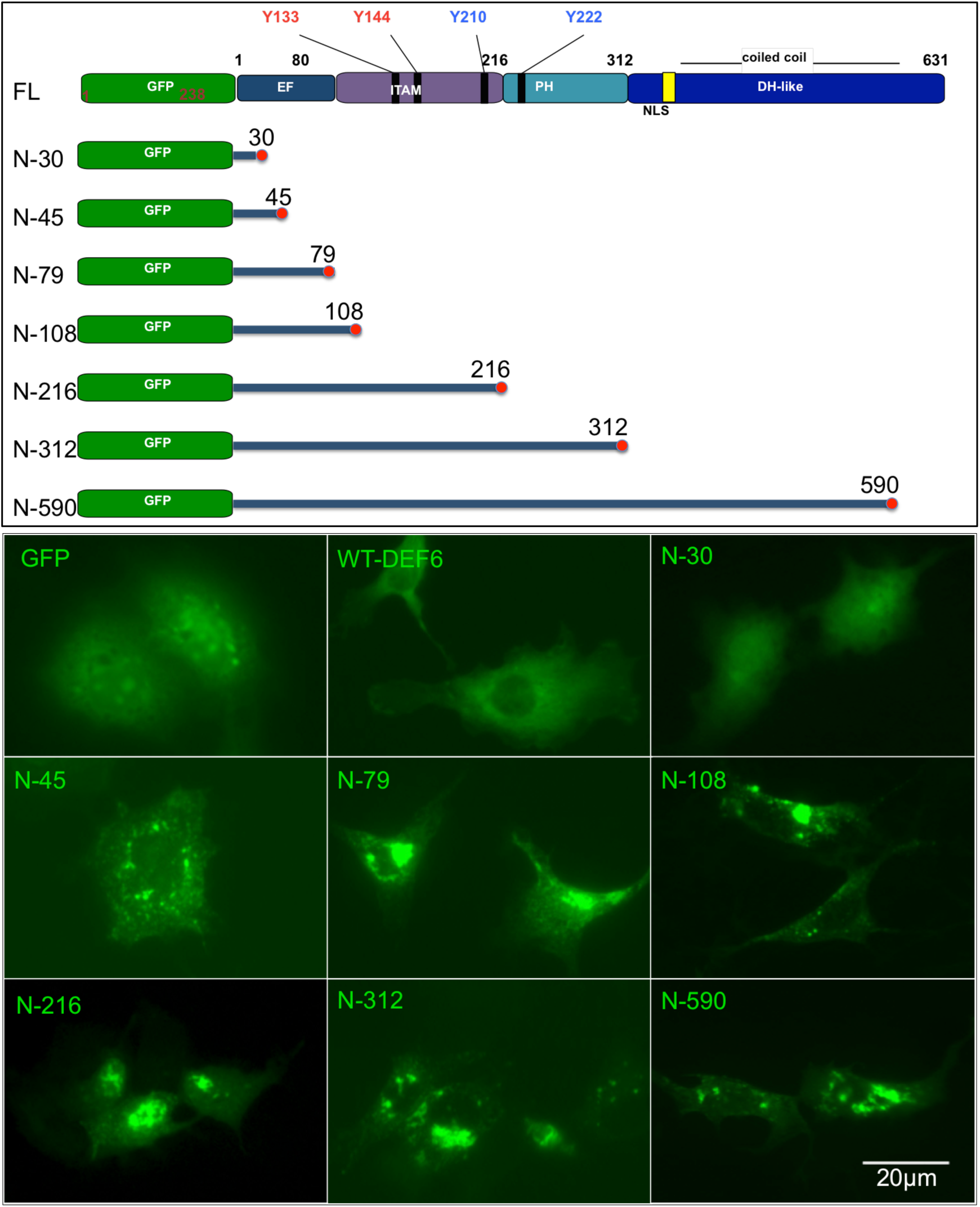
The N-terminal 45 amino acids of DEF6 are necessary and sufficient to form cytoplasmic granules whereas the N-terminal 30 amino acids did not form granules. Upper panel: Schematic representation of C-terminal truncation mutants as indicated. Lower panel: after 24h expression in COS7 cells, GFP-tagged N-45, 79, 108, 216, 312 and 590 exhibited cytoplasmic granules. N-30 however did not form any granules but was diffuse in the cytoplasm and cell nucleus. GFP and wild type DEF6 are shown for comparison.

**Figure 7.**
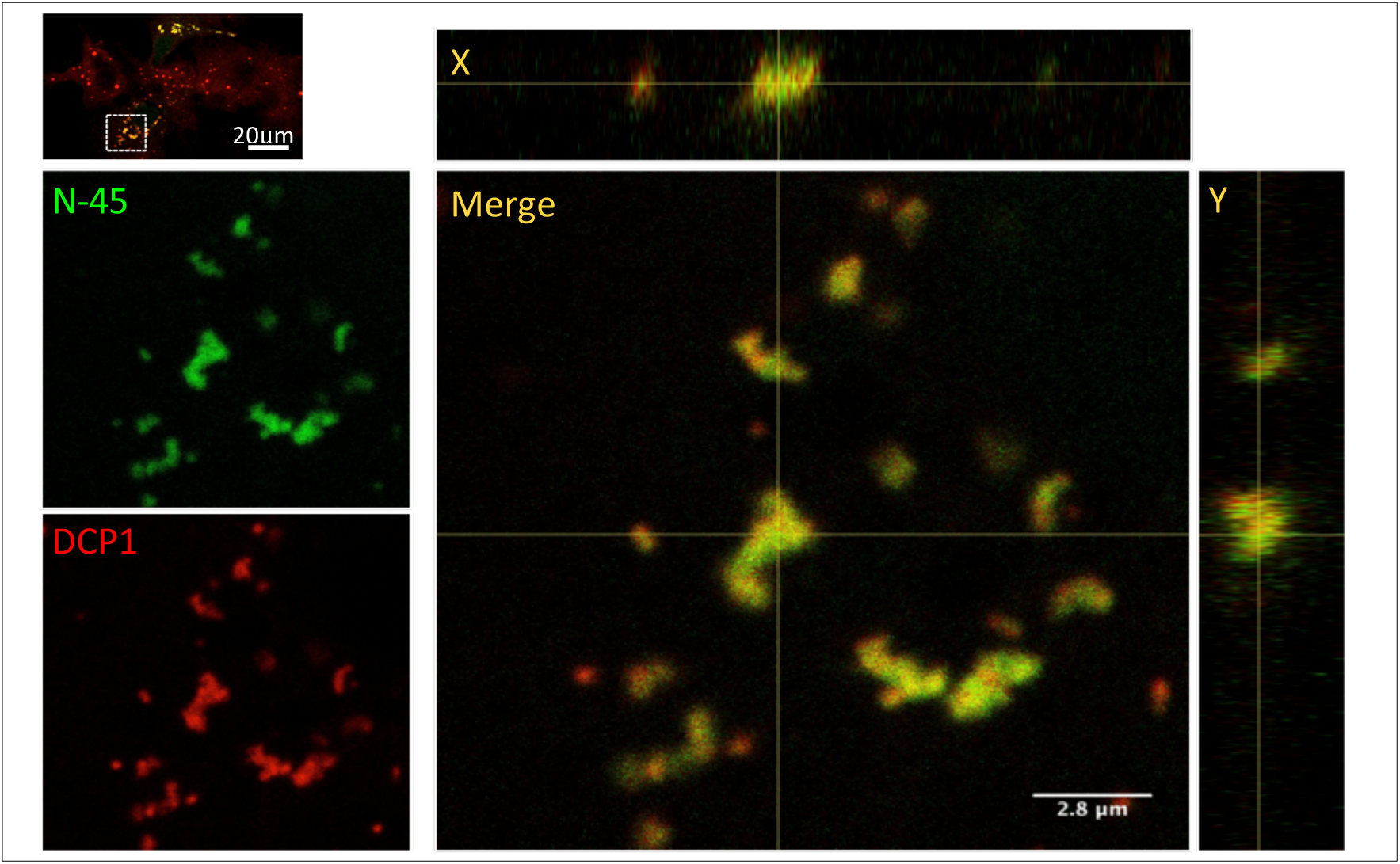
The N-terminal 45 amino acids of DEF6 are sufficient for colocalisation with P-bodies. Confocal Z-stack analysis of GFP-tagged N-45 (green) cotransfected with mCherry-tagged DCP1 (red) 24 h after transfection of COS7 cells. Boxed area in the images on top left are enlarged on the left middle and bottom and further enlarged in the merged images on the right. Vertical and side views (X and Y coordinates) are also shown indicating colocalisation.

## DISCUSSION

To test whether the coiled coil domain is required for granule formation and P-body localisation, proline residues were introduced in highly conserved a/d positions of the heptad repeats to disrupt the coiled coil domain (Fig. 1). Proline mutants did not form granules under cellular stress conditions (Fig. 3) but introduction of Y210/222E into proline mutants resulted in spontaneous granule formation colocalising with DCP1 (Fig. 4). This suggested that the coiled coil domain is dispensable for both granule formation and P-body localisation. Hey et al. (2012) demonstrated that tyrosine (Y) residues 210 and 222 of DEF6 are phosphorylated by ITK in Jurkat T cells and that phosphomimic mutant Y210/222E spontaneously formed cytoplasmic granules that colocalised with P-body marker DCP1 (14). In addition, they showed that treatment of COS7 cells expressing GFP-tagged wild type DEF6 with sodium arsenate resulted in the formation of DEF6 granules colocalising with DCP1. Although COS7 cells do not express ITK, these data suggested that arsenate treatment resulted in modification of DEF6 (perhaps through phosphorylation by an unknown kinase) causing a conformational change that unmasked a region within DEF6 mediating P-body colocalisation. Several C-terminal truncation mutants were generated and tested whether they colocalise with DCP1 when coexpressed in COS7 cells. As shown in Figures 6 and 7, several C-terminal truncated mutants fused to GFP spontaneously formed cytoplasmic granules that colocalised with DCP1 in COS7 cells. Indeed, the N-terminal 45 amino acids were sufficient to target P-bodies whereas localisation of N-terminal 30 amino acids fused to GFP was diffuse in the cytoplasm (N-45; N-30; Fig. 6 and 7). The N-terminal end of DEF6 contains two Ca2+–binding EF hand motifs (26), one of them is present in N-45 but disrupted in N-30. It is not known whether Ca2+-binding is required for P-body colocalisation of N-45, but previous research indicated that Ca2+ signalling influences P-body dynamic. Kilchert et al. (2010) reported that high intracellular Ca2+ promoted P-body formation in yeast and that the P-body component ALG-2, that has five EF hand motifs, capable of binding Ca2+, interacts with PATL-1 is in a Ca2+-dependent manner. The first Ca2+-binding motif of DEF6 is located within the N-terminal 45 amino acids and disruption of it abolishes P-body colocalisation. It is therefore tempting to speculate that Ca2+ mediated signalling is linked to DEF6 P-body dynamics. However, in an accompanying work (16), we have shown that under cellular stress conditions, the N-terminal end of DEF6 is dispensable for granule formation and P-body colocalisation suggesting that other signals might also be able to direct C-terminal domains of DEF6 to P-bodies. The function of DEF6 in P-bodies is not know, but given that N- and C-terminal domains of DEF6 are involved in P-body colocalisation indicates the complexity of the molecular mechanisms involved and perhaps implying that DEF6 has multiple roles in mRNA metabolism in P-bodies.

## EXPERIMENTAL PROCEDURES

COS7 cells were cultured in 10% (v/v) FBS (Sigma, #D6429) supplemented DMEM (Sigma, #D6429) at 37°C, 5% C02 and humidified atmosphere. The original eGFP-C2-DEF6 plasmid (5) was used to generate the mutant DEF6 proteins by mutagenesis PCR (QuikChange Lightning Multi Site-Directed Mutagenesis Kit, Agilent Technologies, #210516). All mutations were verified by sequence analysis (see Suppl. Fig, 4 and 5). WT and mutated DEF6 plasmids were transfected (Gene-juice Transfection Reagent, Novagen, #70967-3) into COS7 cells and 24h after the transfection, cells were fixed with 4% PFA at room temperature and analysed using ZEISS LSM 710 confocal microscope. Image analysis was performed using Fiji software. mCherry-tagged DCP1 plasmid was applied as P-body marker premixed 1:1 (w/w) with WT or mutant DEF6 plasmids for co-transfection. Arsenate (1mM, Sigma 50mM stock solution, #35000) treatment of transfected cells was carried out at 37°C for 40 mins before fixation of the cells.

## Acknowledgments

We thank Drs. Martin Gering, Sally Wheatley and Peter Jones for helpful discussions, the SLIM team at the University of Nottingham for their support in confocal microscopy and the undergraduate students Michael Darrington and David Chandler for their assistants in establishing some of the DEF6 mutants.

## FOOTNOTES

The abbreviations used are: DEF6, Differentially Expressed in FDCP 6; IBP, IRF4-binding protein; SLAT, SWAP-70-like adaptor protein of T cells; IS, immunological synapse; P-body, mRNA processing body; DCP1, decaping protein 1

