## Supplementary material for "N-terminal 45 amino acids of DEF6 are necessary and sufficient to spontaneously colocalise with DCP1 in P-bodies": Suppl.

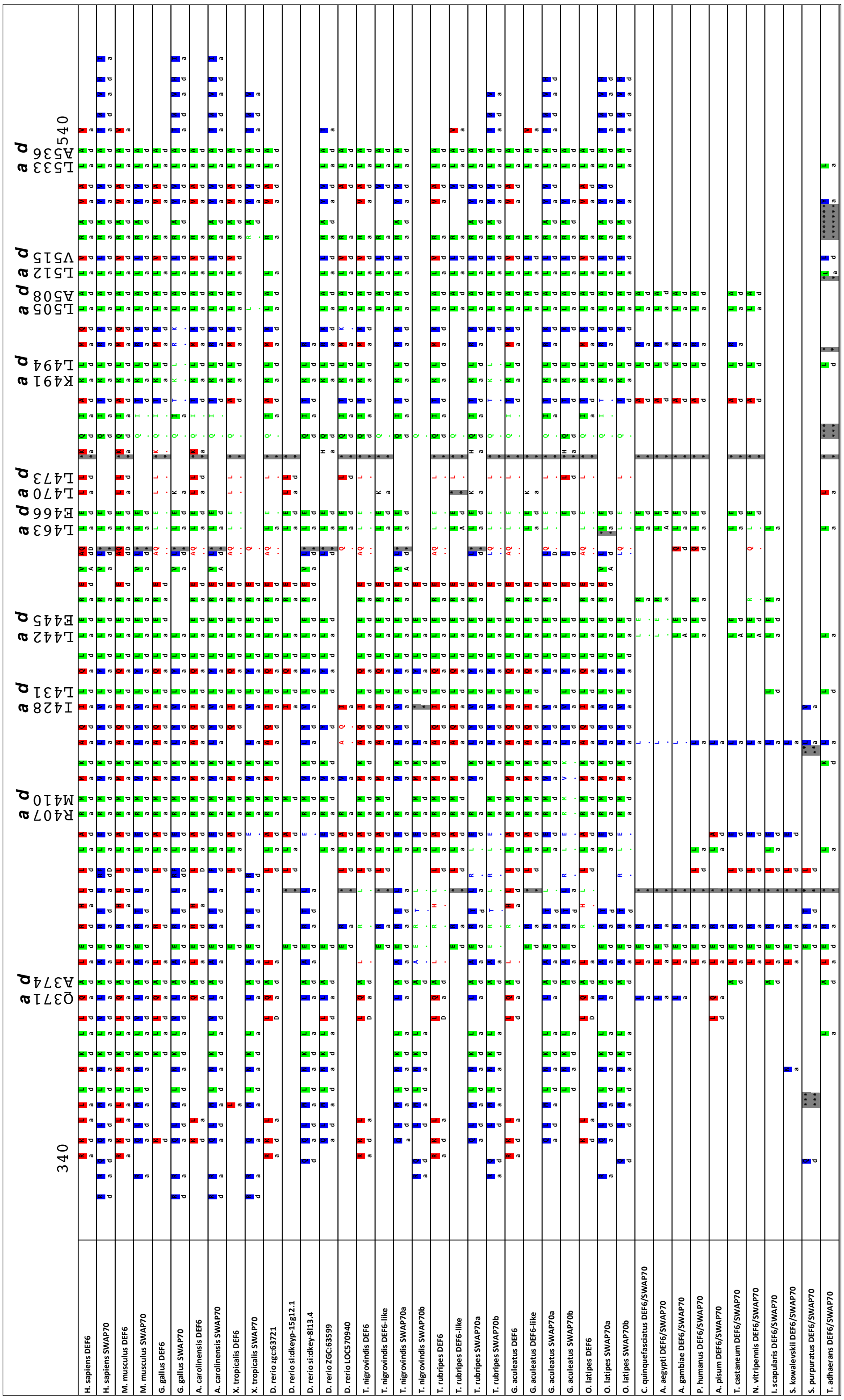

**Supplementary Figure 1.** Comparison of the "a" and "d" positions within the coiled coil domain of DEF6 and SWAP70 across species from human to trichoplax. Colour coding of amino acid residues indicates identity: red – amino acids identical to the ones in human DEF6; blue – amino acid identical to the ones in human SWAP70; green -amino acids indentical to both human DEF6 and SWAP70 (Spencer and Sablitzky, unpublished).

```
1        10        20        30        40        50        60
MALRKELLKSIWYAFTALDVEKSGKVSKSQLKVLSHNLYTVLHIPHDPVALEEHFRDDDD
          ________////////////________          _______////
           helix  Ca2+-binding helix             helix  Ca2+

61       70        80        90        100       110       120
GPVSSQGYMPYLNKYILDKVEEGAFVKEHFDELCWTLTAKKNYRADSNGNSMLSNQDAFR
////////_______
-binding  helix
```

**Supplementary Figure 2.** N-terminal amino acid sequence of DEF6. DEF6 N-terminal contains two Ca2+-binding motifs, from amino acid 10 to 40, and 50 to 75.

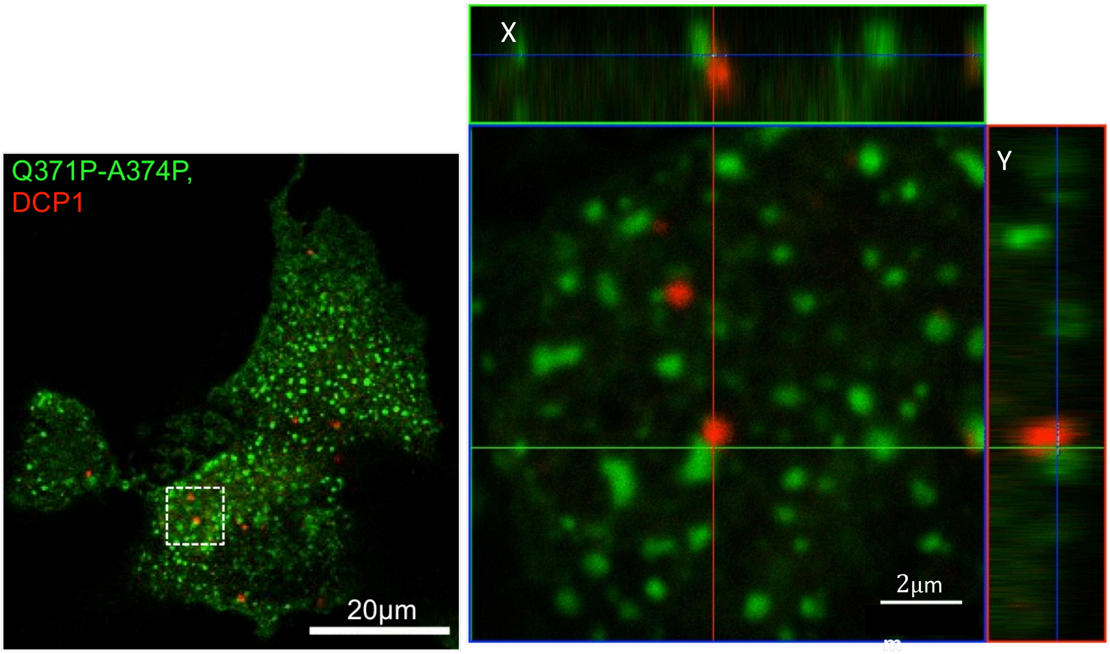


**Supplementary Figure 3.** Q371P-A374P mutant granules are not colocalising with P-bodies. Confocal Z-stack image analysis of Q371P-A374P (green) and P-body marker DCP1 tagged with mCherry (red) cotransfected in COS7 cells. Boxed area in the left image is shown enlarged on the right. Vertical and side views (X and Y coordinates) are also shown indicating that Q371P-A374P did not colocalise with DCP1.

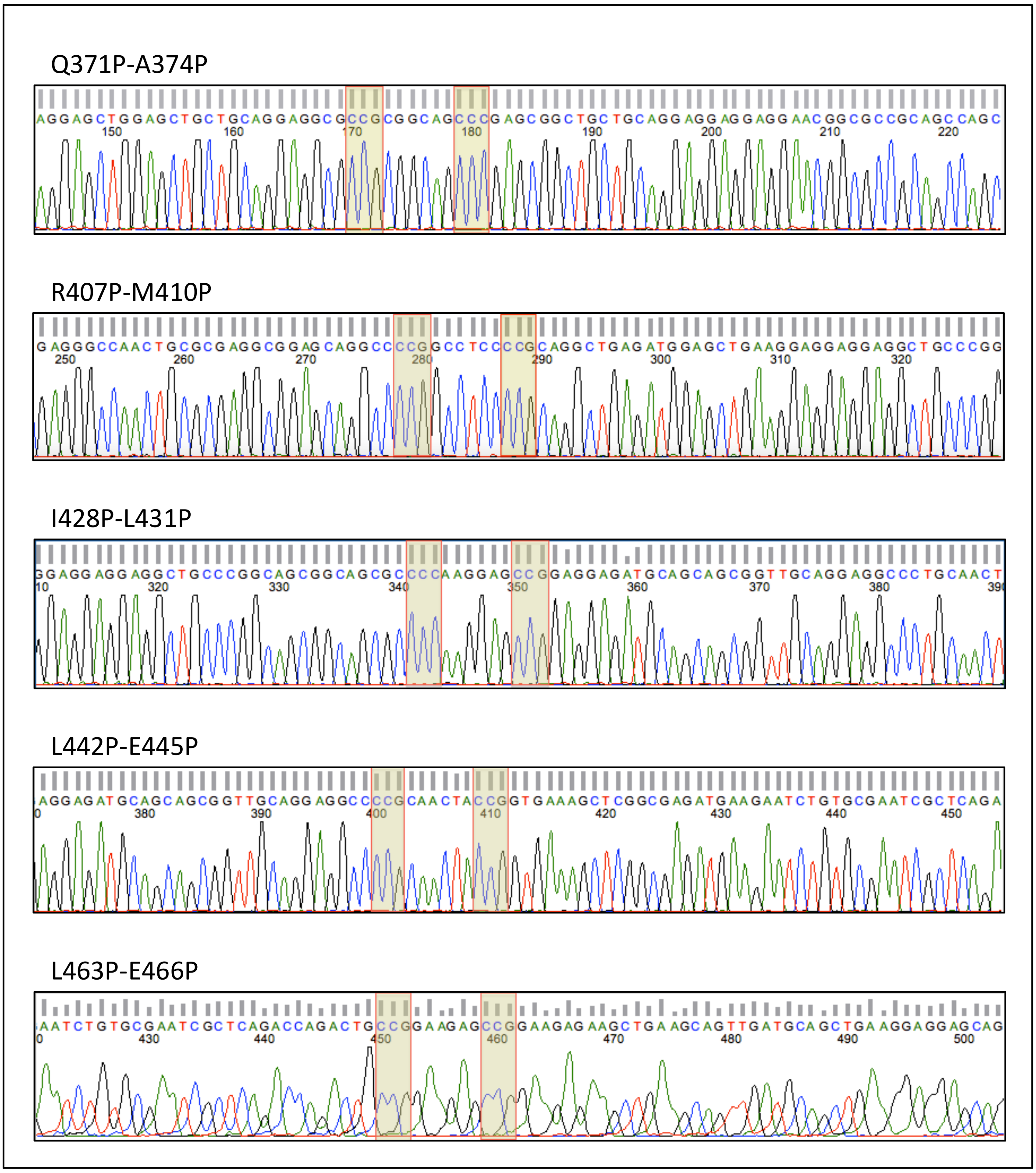


**Supplementary Figure 4.** DNA Sequencing results of DEF6 Proline mutants. The mutated positions are highlighted by coloured boxes. Continued in next three pages.

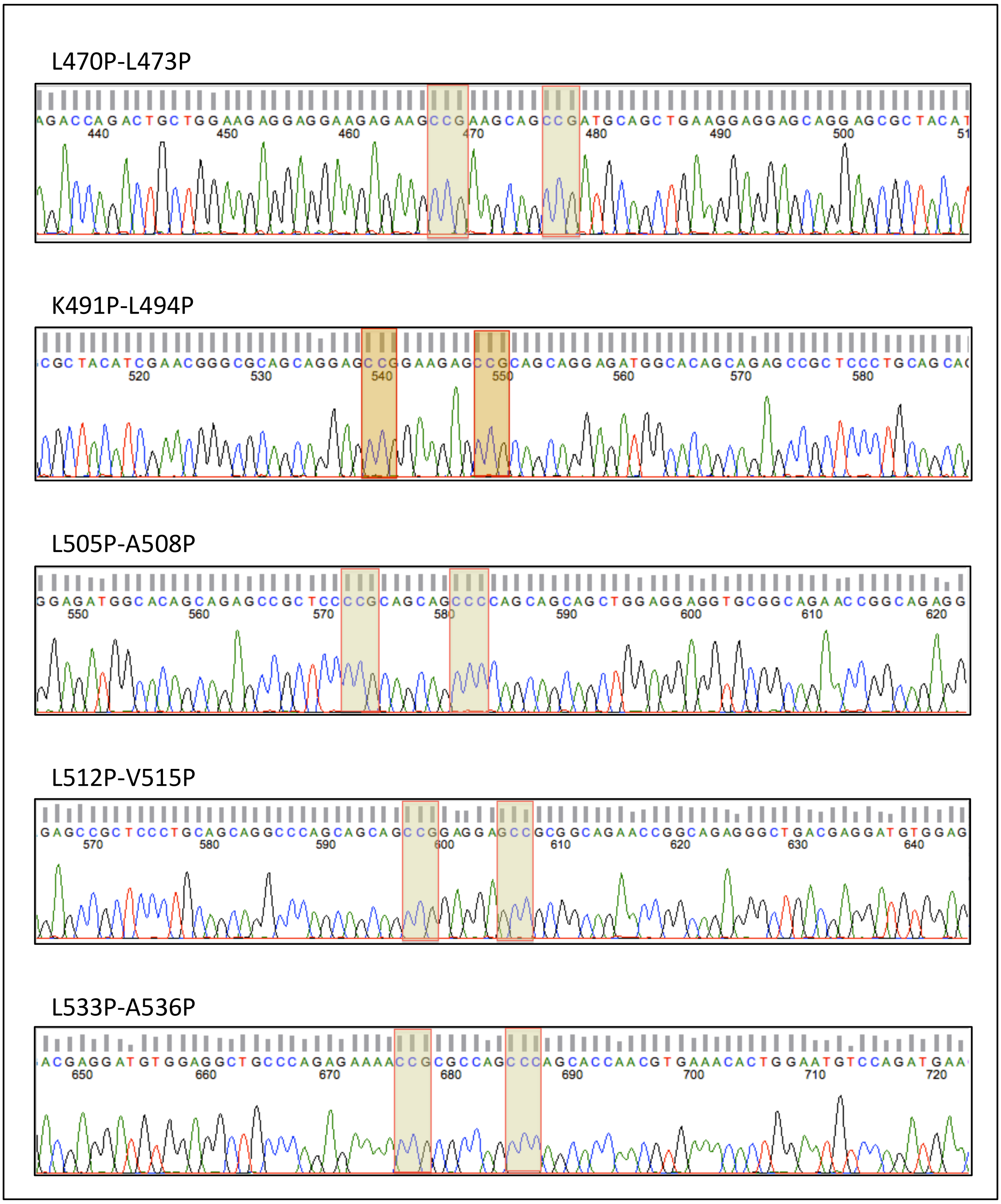


**Supplementary Figure 4.** DNA Sequencing results of DEF6 Proline mutants. The mutated positions are highlighted by coloured boxes. Continued in next two pages.

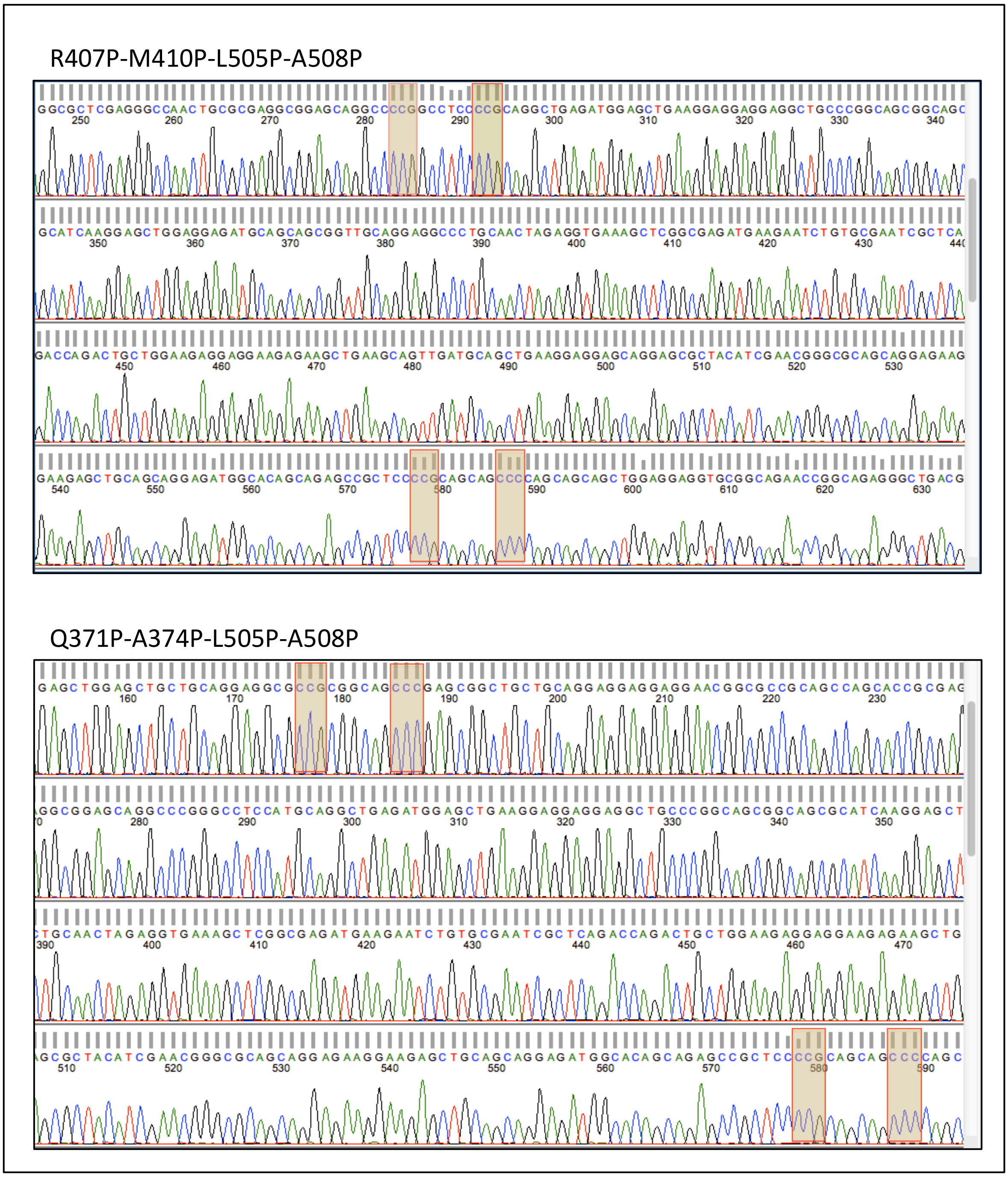


**Supplementary Figure 4.** DNA Sequencing results of DEF6 Proline mutants. The mutated positions are highlighted by coloured boxes. Continued in next pages.

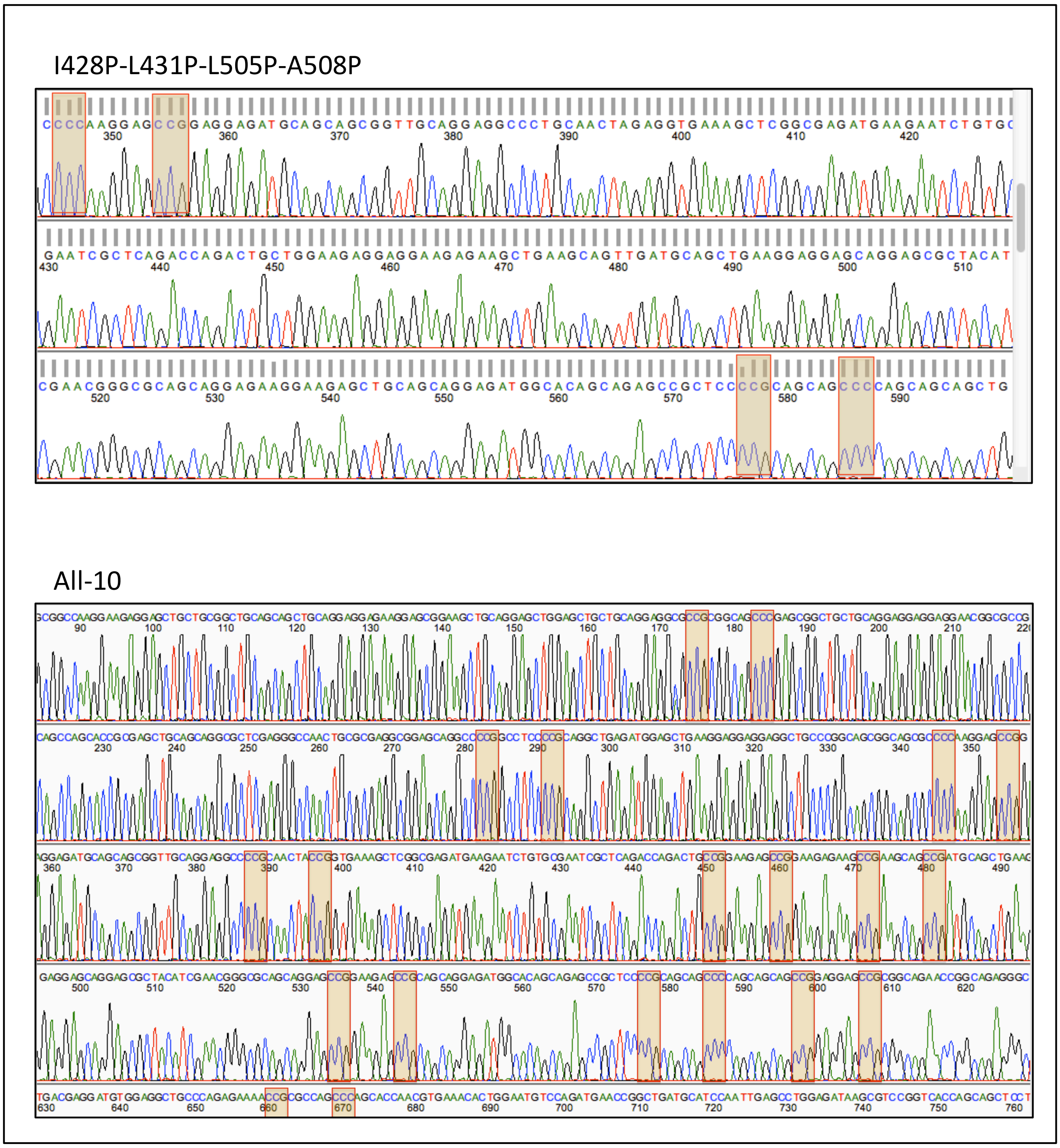


**Supplementary Figure 4.** DNA Sequencing results of DEF6 Proline mutants. The mutated positions are highlighted by coloured boxes

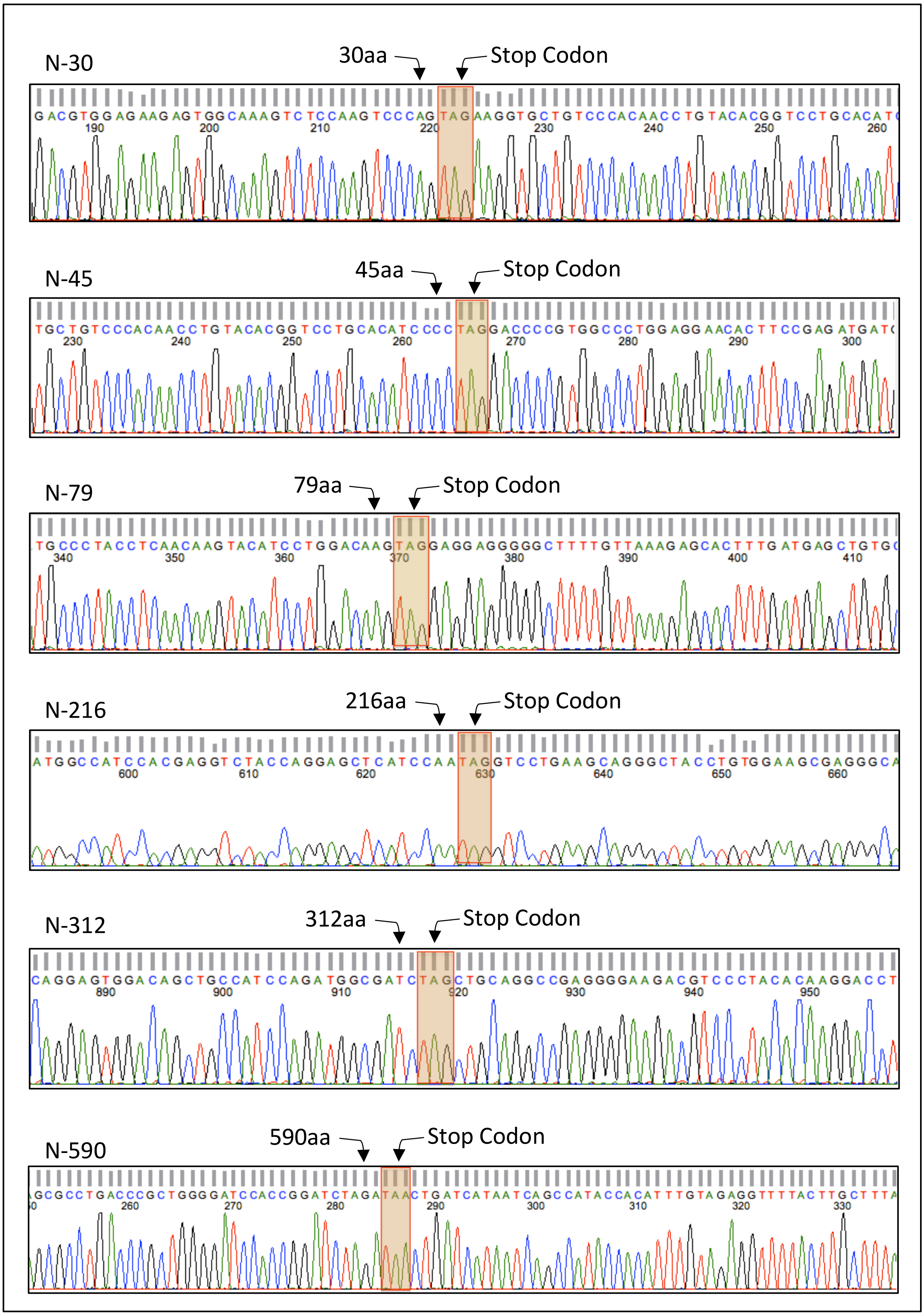


**Supplementary Figure 5.** DNA sequencing results of DEF6 C-terminal truncated mutants. The inserted stop codon and their positions are indicated in coloured boxes.
